# OverFlap PCR – a reliable approach for generation of plasmid DNA libraries containing random sequences without template bias

**DOI:** 10.1101/2022.01.11.475799

**Authors:** Artis Linars, Ivars Silamikelis, Dita Gudra, Ance Roga, Davids Fridmanis

## Abstract

Over the decades the improvement of naturally occurring proteins and creation of novel ones has been the primary goal for many practical biotechnology researchers and it is widely recognized that randomization of protein sequences coupled to various effect screening methodologies is one of the most powerful techniques for fast, efficient and purposeful approach for acquisition of desired improvements. Over the years considerable advancements have been made in this field, however development of PCR based or template guided methodologies has been hampered by the resulting template sequence bias. In this article we present novel whole plasmid amplification based approach, which we named OverFlap PCR, for randomization of virtually any region of the plasmid DNA, without introduction of mentioned bias.

## INTRODUCTION

Ever since the emergence of recombinant DNA technologies and their employment in practical biotechnology for production of desired biological compounds (around the mid-1970s) [1, 2] the improvement of employed proteins and creation of new ones has been considered as the Holy Grail by many researchers working in this field and the first attempts to work towards this goal that were undertaken in early 1980s hallmarked the formation of the fields of protein engineering and synthetic biology [3, 4]. Therefore it is not surprising that over the following years a number of researchers focussed their attention on development and employment of novel strategies and methods for fast, efficient and purposeful alteration of DNA sequences that encode their protein of interest and their insertion into appropriate expression vector. As one would expect, nowadays these efforts have resulted in a multitude of approaches that are still being continuously modified and improved. Although the list of the developed methods is extensive and might be confusing to any newcomer to the field, there are two questions that every researcher must answer before selection of appropriate approach: 1) “Is it necessary to alter protein coding sequence at specific site or at multiple sites?” and 2) “Is it necessary to replace selected codon with specific of random sequence?”.

The site and residue specific alterations represent the group of simplest modifications, thus basic PCR [3], megaprimer [5] or whole plasmid [6] based site-directed mutagenesis coupled to restriction - ligation [1], Gibson assembly [7], recombination [8] or deoxy uridine - USER enzyme (Uracil-Specific Excision Reagent) [9] based cloning into selected vector are usually employed. All these methods are reliable and proven to provide excellent results in multitude of publications. The introduction of residue specific alterations at multiple sites however is more challenging. The simplest of solutions would involve multiple repetitions of single site directed mutagenesis cycles, but this approach is time and labour consuming. Therefore such methods as Progressive PCR Based Multi-Site-Directed Mutagenesis [10], Mutant Strand Synthesis By Primer Extension And Ligation [11], Homologous Recombination Based Multi-Site-Directed Mutagenesis [12], Multichange Isothermal (MISO) Mutagenesis [13] Quikchange Multiple Site-Directed Mutagenesis [14] and many others, including various modifications of already mentioned, [15-20] where developed. Despite this great variety in approaches all these methods are just as reliable as single site mutagenesis methods, because the success in both cases is determined by the necessity to acquire single clone with desired alterations.

The success for randomization approaches on the other hand is determined by the ability of method to deliver the library of sequences with the desired level of diversity and greatest possible number of variants. Thus the selection of appropriate method and evaluation of its limitations is of paramount importance. When considering multi-site randomization, four types of approaches that differ significantly in type of acquired sequence libraries can be distinguished: 1) random mutations at random positions – these are usually introduced through employment of Error-Prone PCR [21-26] or E.coli mutator strains [27-30]; 2) random combinations of multiple site and residue specific alterations – introduced with the help of Split-mix-PCR [31], RECODE [32], OSCARR [33] or several others; 3) random recombination of multiple pre-existing DNA sequences – acquired through DNA shuffling [34-36], StEP [37, 38], RACHITT [39, 40] or ITCHY [41-44]; and 4) random residues at multiple specific positions - usually introduced in similar manner as residue specific alterations at multiple sites with the exception that, instead of specific alteration containing oligonucleotides, a library of oligonucleotides that randomize selected position are employed [45]. According to literature data the methods for the first three are either already well developed and are used for many years without significant alterations (error prone replication/amplification) or are constantly being redesigned at their core providing libraries of greater quality or procedure of simpler design (other two). However, in regard to randomization of residues at multiple specific positions, as of recent significant developments have not been made, because those single site methods that serve as the basis for these approaches are sufficiently effective and reliable for introduction of small – up to 6 nucleotide (2 aa codon) randomizations at single site [46], which in essence is the target number for majority of studies. The necessity for introduction of larger scale randomizations is also recognized, as it would be beneficial for examination of larger protein motifs, but it is rarely employed, because it usually results in library with significant sequence distribution bias towards the employed template. The only suggested means for reduction of this effect that we encountered were careful design of the oligonucleotides [47] and fine tuning of oligonucleotide annealing temperature [48], which is also recommended if 6 or less nucleotides are randomized.

Consequentially, this bias problem is also acute in the case of single site randomization, because the same methods that are used for site and residue specific alterations are also employed here [23, 47, 49, 50]. As one can see from information that was provided earlier, majority of these methods are PCR or other type of template based, thus also in here the if the target motif is larger than few codons then oligonucleotides with the greatest similarity to template shall bind/anneal with greater efficiency than others and the whole process shall result in acquisition of library with decreased sequence diversity. However unlike in residue specific replacement here are several additional frequently used methods, which were specifically designed to minimize this bias. The most prominent of these is the cassette mutagenesis [51-53] where restriction/ligation is used to insert chemically synthesized randomized sequence containing fragment into the target site. However despite its apparent advantages this method has a few technical drawbacks. As of first there is the apparent need of conveniently located restriction site/s and if such are not available these have to be introduced using site directed mutagenesis and without (or minimal) disruption of encoded aa sequence. As of second, in our experience the efficiency of two fragment ligation is significantly lower than that of circularization, thus its employment limits the diversity of acquired library and Neme *et al*. [53] reported that employing this strategy ∼1000 unique variants per experiment were reliably identified. Loop-out/loop-in is another method that addresses the bias issue [54]. Although this method is classical PCR mutagenesis based, here the control over the undesired selectivity is exerted though excision of target site during the first round of mutagenesis and insertion of randomized sequence during the second. This approach indeed should eliminate the bias towards the target site, but it might introduce the bias towards the adjacent and hairpin structure forming sequences and since this method is infrequently employed the literature data on its drawbacks is scarce.

Another aspect that should nowadays be considered when working with randomized libraries is their quantitative characterization for both quality assessment and experimental purposes (acquired data might serve as point 0 in experiments where changes in relative abundancy of clones are assessed). For the studies that were carried out more than a decade ago Sanger sequencing was pretty much the only available option, therefore involved personnel was bound to hand pick individual colonies to acquire sequence of inserted fragment. Since such strategy is laborious, costly and low throughput, it usually resulted in acquisition of low data amounts, but nowadays with the widespread availability of massive parallel sequencing technologies (also known as NGS) this limitation is largely overcome [15, 53, 55, 56]. However, surprisingly, we also observed that in a number of reports particularly in those that claimed development of “novel and highly effective” randomization strategy NGS data was not presented [49, 57], which indicates that applicability of presented methodology to selected tasks should be reviewed critically and presented claims considered with caution.

Thus in the light of presented information we believe that in the case of specific multi- and single-site mutagenesis the methods for creation of random sequences libraries without template bias are still insufficiently developed and there is a room for significant improvements. The underlying purpose of this methodological study was to create an expression plasmid library for secreted production of 18 aa long random peptides in yeast *Saccharomyces cerevisiae*, which shall further be used for screening of novel biologically active molecules. During the course of these activities and while struggling with the described bias we developed a novel approach for reliable introduction of random nucleotide sequence within virtually any site of the plasmid, which we named the “OverFlap PCR” to emphasize the distinction from “Overhang PCR” and “Overlap PCR”. During the preparation of this article intense internal discussions were started over the possibilities of further methodological improvements of method, which shall also be presented in subsequent sections.

## MATERIALS AND METHODS

### Creation of *S. cerevisiae* Compatible Secreted Peptide Expression Plasmid p426GPD-αfactor-αMSH

To enable the secretion of produced peptides p426GPD expression plasmid, kindly provided by Dr. Simon Dowell from GSK (Stevenage, UK), was supplemented with coding sequence (CDS) for *S*.*cerevisiae* α-factor secretion signal. To create the source plasmid that could be used as positive control during expression experiments a CDS of α-Melanocyte stimulating hormone (α-MSH) was added to 3’ end of secretion signal’s CDS. Both elements were inserted simultaneously in following manner.

CDS of secretion signal was amplified by PCR from 0.05 μg of pPIC9K vector plasmid (Thermo Fisher Scientific, USA) using 2.5 U of Pfu polymerase (Thermo Fisher Scientific, Lithuania), 10 pmol of aFactor-BamHI-Fw forward primer (all primers used in this study were purchased from Metabion GmbH, Germany), which contained *Bam*HI restriction site and 10 pmol of aFactor-aMSH-EcoRI-Rs reverse primer, which contained *Eco*RI restriction site and α-MSH CDS (Table 1). Other reaction reagents included 2μl of 10x reaction buffer with MgSO_4_ (Thermo Fisher Scientific, Lithuania), 4 nmol of each dNTP (Thermo Fisher Scientific, Lithuania) and water to final volume of 20μl. The reaction was performed in Veriti PCR thermal cycler (Applied Biosystems, USA) under following conditions 95 °C for 5 min; followed by 40 cycles of 95 °C - 15 sec, 63 °C - 30 sec, 72 °C – 60 sec and finalized with 72 °C - 5 min. The verification of success and purification of acquired product from template and primer dimers was performed by preparative agarose gel electrophoresis, appropriate band excision, purification employing GeneJET Gel Extraction Kit (Thermo Fisher Scientific, Lithuania) and elution in 18 μl of ultrapure water. Then both, whole volume of purified PCR product (αFactor-αMSH) and 2 μg of p426GPD vector plasmid, where cleaved with *Bam*HI (Thermo Fisher Scientific, Lithuania) and *Eco*RI (Thermo Fisher Scientific, Lithuania) restriction enzymes in BamHI reaction buffer and 20μl of total reaction volume according to manufacturer’s instructions. Following the restriction both fragments (fαFactor-αSHM and v426GPD) were purified using GeneJET PCR Purification Kit (Thermo Fisher Scientific, Lithuania) according to manufacturer’s instructions and eluted in 20 μl of ultrapure water. After that 1μl of v426GPD and 7 μl of fαFactor-αMSH were mixed with 1 μl of 10x T4 DNA ligase reaction buffer and 5U of T4 DNA ligase (Thermo Fisher Scientific, Lithuania), incubated for 1 h at 22°C and whole reaction volume was transformed in chemically competent *E*.*coli* Dh5α strain cells (acquired from Invitogen, USA and prepared according to Green & Rogers instructions [58]), which were then seeded onto ampicillin supplemented LB media petri dish and incubated overnight at 37°C. The success of fragment insertion was verified through agarose gel electrophoresis visualization of colony PCR products (1xTaq reaction buffer; 17.5 nmol of MgCl_2_; 4 nmol of each dNTP; 10 pmol of each M13-Fw and M13-Rs primer (Table 1); 0.5 U of recombinant Taq DNA Polymerase (Thermo Fisher Scientific, Lithuania) and water to final volume of 10μl; thermal conditions: 95°C for 5 min; followed by 45 cycles of 95 °C - 30 sec, 55 °C - 30 sec, 72 °C – 1 min 30 sec and finalized with 72 °C - 7 min). Colonies that produced DNA fragment of 1300 bp length were considered as insertion positive. Two of the positive colonies were inoculated in 6 ml of 2YT media, incubated overnight at 37°C in shaker and the plasmid DNA was extracted using GeneJET Plasmid Miniprep Kit (Thermo Fisher Scientific, Lithuania). As the final means of verification sequence of inserted fragment was verified through Sanger Sequencing employing M13-Fw and M13-Rs primers, BigDye™ Terminator v3.1 Cycle Sequencing Kit (Thermo Fisher Scientific, USA), according to manufacturer’s instructions and 3130/3130xl Genetic Analyzer (Applied Biosystems, USA). The selection of plasmid for further activities was based on quality of acquired DNA sequences i.e. only the clones that yielded high quality chromatograms were employed. Map of thus acquired plasmid is presented in Supplementary Figure 1.

**Table 1.**
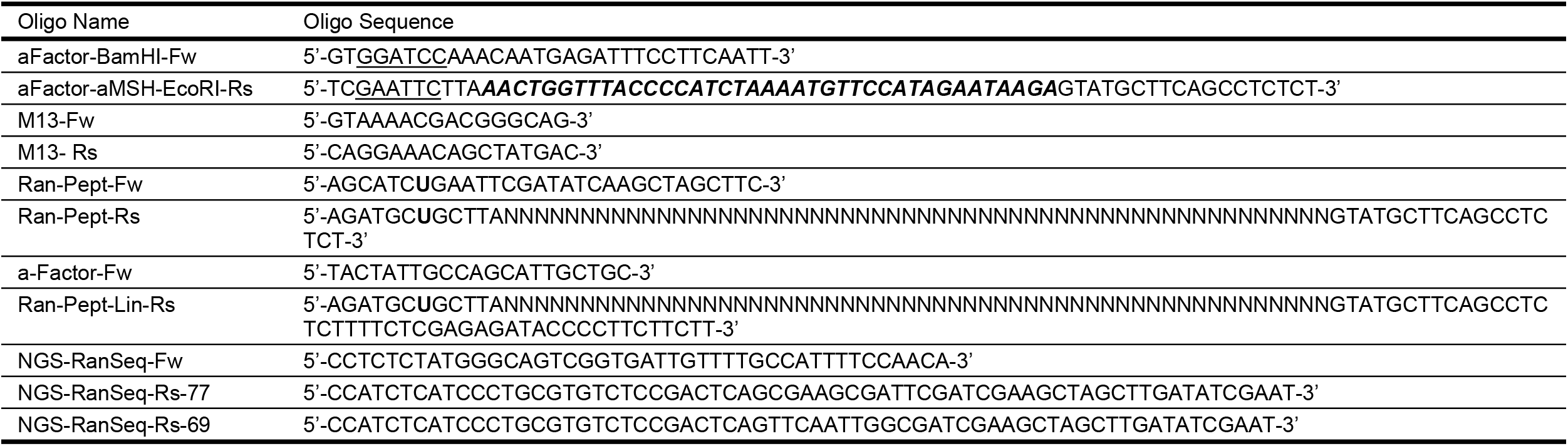
Employed oligonucleotides. Restriction sites are in <u>underline</u>, coding sequences are in ***Bold-Italic*** and deoxy uridine in **Bold**

### Randomization of peptide CDS

#### Randomization employing modified whole plasmid amplification strategy (WPA)

Our initial approach for randomization of peptide coding sequence was based on modified whole plasmid amplification strategy [6, 50], the modifications included employment of deoxy uridine containing primers and USER enzyme mix (New England Biolabs, UK) to create sticky ends that would facilitate the circularization of plasmid [9]. Principal scheme of the procedure is presented in Figure 1.

**Figure 1.**
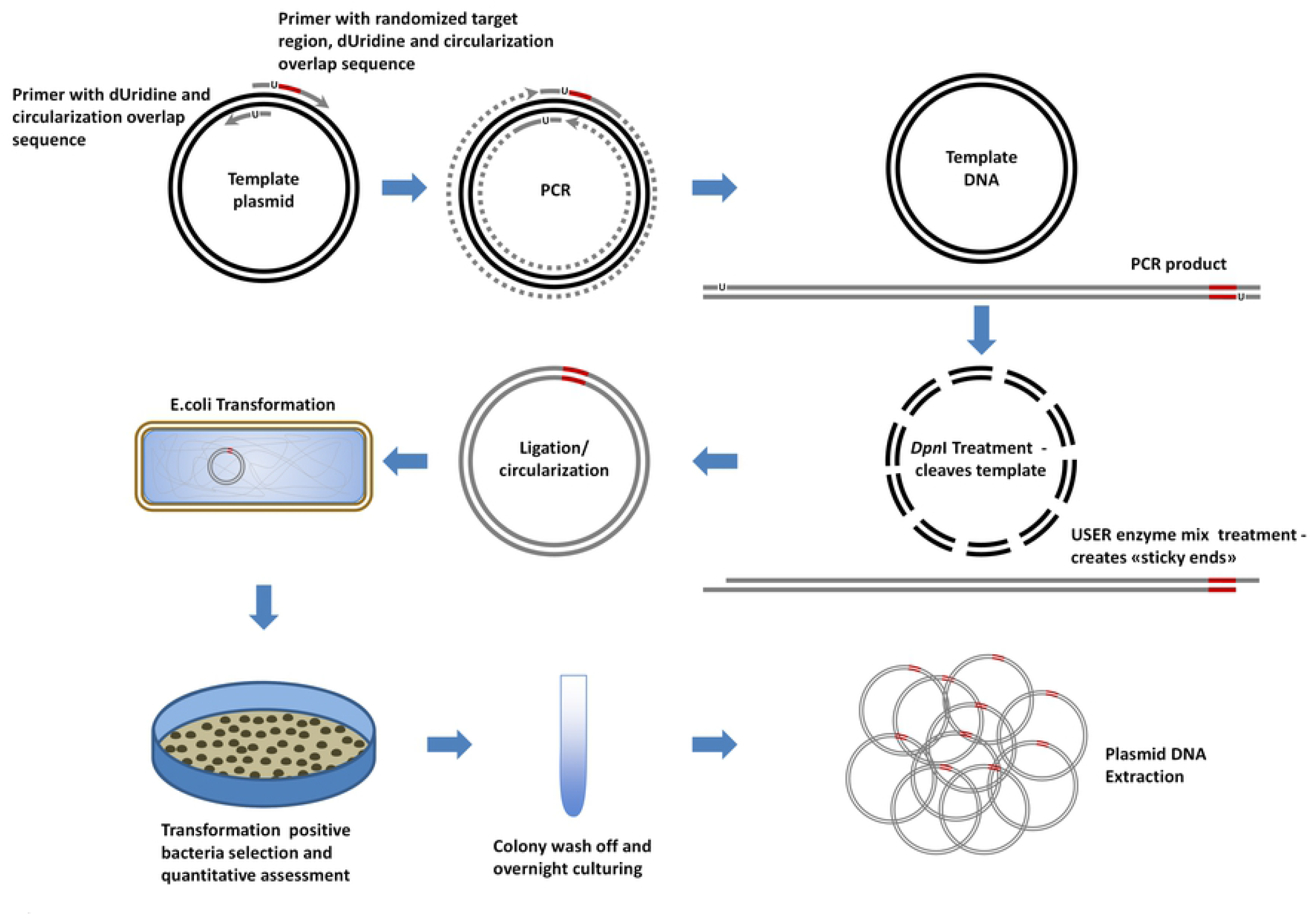
The principal scheme of employed strategy for whole plasmid amplification based randomization of selected DNA region. Randomization is carried out employing modified whole plasmid amplification strategy (WPA), where we employed 1) one long reverse strand oligonucleotide, which starting from 3’ end contained sequence that is complementary to plasmid DNA, randomized region (represented by red line), stop codon and additional sequence where the first thymidine was replaced with deoxy uridine (represented by U), 2) one short forward strand oligonucleotide, which starting from 3’ end contains sequence that is complementary to plasmid DNA and sequence that is complementary to reverse primer’s 5’ nucleotides (here also the first thymidine was replaced with deoxy uridine), and 3) high fidelity polymerase (Phusion U) which tolerates presence of Uridine. Following the amplification, reaction mixture was treated with *DpnI* restriction enzyme, which cleaves only methylated (bacterial origin) DNA, and USER (Uracil-Specific Excision Reagent) enzyme mix, which excises uridine base from amplification product and cleaves abasic site, thus forming “sticky end”. The amplification product is then circularized and transformed into competent *E*.*coli* Dh5α strain cells, which are then seeded onto ampicillin supplemented LB media petri dish and incubated overnight at 37°C. The next day transformation positive colonies were quantified, washed off, inoculated in ampicillin containing liquid media and following overnight culturing employed in extraction of Randomized Plasmid DNA. The synthesized chain is represented by grey line; Dotted line represents DNA synthesis; and dashed line represents template DNA degradation by DpnI restriction endonuclease.

The whole plasmid amplification strategy (WPA) reaction mixture contained 1x Phusion U Multiplex PCR Master Mix (Thermo Fisher Scientific, Lithuania), 0.06 μg of p426GPD-αfactor-αMSH plasmid, 50 pmol of Ran-Pept-Rs primer, which in essence was identical to previously employed aFactor-aMSH-EcoRI-Rs primer, but with α-MSH CDS replaced with 54 random nucleotides and at 5’ end containing artificial sequence of 6 GC50% nucleotides that end with uridine (Table 1), 50 pmol of Ran-Pept-Fw primer, which staring from 5’ end contains 6 nucleotides that are complimentary to reverse primer’s artificial GC50% sequence and 23 nucleotides that are complementary to sequence that follows immediately after α-MSH CDS of template plasmid, and water to final volume of 50μl; thermal conditions were as follows: 98°C for 3 min; followed by 10 cycles of 98 °C - 10 sec, 59 °C - 30 sec, 72 °C – 3 min 30 sec and finalized with additional 72 °C – 7 min. Success of the reaction was verified by agarose gel electrophoresis and acquired products were for 1 h at 37°C simultaneously treated with 2 U of USER enzyme mix and 20 U of *Dpn*I restriction enzyme (Thermo Fisher Scientific, Lithuania). The first reagent introduces nicks at the uridine site thus releasing first six 5’ nucleotides and creating sticky end, while DpnI exclusively cleaves methylated recognition sites, which are typically found only in DNA that has been extracted from bacteria, thus in essence shearing only the template DNA. DNA fragments were then purified using GeneJET PCR Purification Kit, eluted in 20 μl of ultrapure water and 8 μl of purified DNA were used for circularization employing T4 DNA ligase as described earlier, subsequent transformation of whole reaction volume in chemically competent *E*.*coli* Dh5α strain cells (prepared inhouse employing “Rubidium chloride competent cell protocol” which is available at https://mcmanuslab.ucsf.edu/protocol/rubidium-chloride-competent-cell-protocol and estimated transformation efficiency of 109 CFU per 1μg of pUC19 plasmid), which were then seeded onto ampicillin supplemented LB media petri dish and incubated overnight at 37°C. The following day number of transformation positive colonies (and possible randomization clones) was estimated by petri dish imaging in UVP Biospectrum® AC Imaging System (UVP, USA) and colony count estimation by OpenCFU 3.9.0 software [59] (available at http://opencfu.sourceforge.net/). Following that all colonies were washed off with 1 ml and inoculated in 5 ml of ampicillin supplemented 2YT media and incubated overnight at 37°C in shaker. The plasmid DNA extraction and sequencing was performed as described previously, only in this case a-Factor-Fw (Table 1) was used instead of M13-Rs primer.

#### Randomization employing OverFlap PCR strategy (OverFlapWPA and OverFlapAsymWPA)

0.2 μg of created p426GPD-αfactor-αMSH plasmid where cleaved with *Xho*I restriction enzyme (Thermo Fisher Scientific, Lithuania) in R reaction buffer and 20μl of total reaction volume according to manufacturer’s instructions. The success of the reaction was verified by agarose gel electrophoresis and resulting fragment was purified employing GeneJET PCR Purification Kit. Further 0.06 μg of linearized plasmid were used for whole plasmid amplification as described previously, with two exceptions – 1) in one of reactions asymmetric amplification (Asym) was intended during initial cycles to avoid exponential amplification of initial variants (OverFlapAsymWPA) therefore forward primer was not added and total reaction volume was 45 μl and 2) randomization for both reactions (OverFlapWPA and OverFlapAsymWPA) was performed employing Ran-Pept-Lin-Rs primer (Table 1), which unlike previously employed Ran-Pept-Rs primer contained additional sequence of 26 nucleotides that is complementary to α-factor secretion signal CDS’s 3’ end, thus covering the 18 nucleotide sequence upstream of *Xho*I restriction site. Thermal conditions for reaction with both primers (OverFlapWPA) was as described previously while for the other one (OverFlapAsymWPA) they were as follows: 98°C for 3 min; followed by 10 cycles of 98 °C - 10 sec, 59 °C - 30 sec, 72 °C – 3 min 30 sec and finalized with 98 °C - 3 min. During the last incubation 50 pmol of Ran-Pept-Fw primer were added directly to reaction mixture and reaction was continued for additional 25 cycles, which were finalized with additional 72 °C – 7 min. All following activities were done in identical manner as previously. Principal scheme of the procedure is presented in Figure 2.

**Figure 2.**
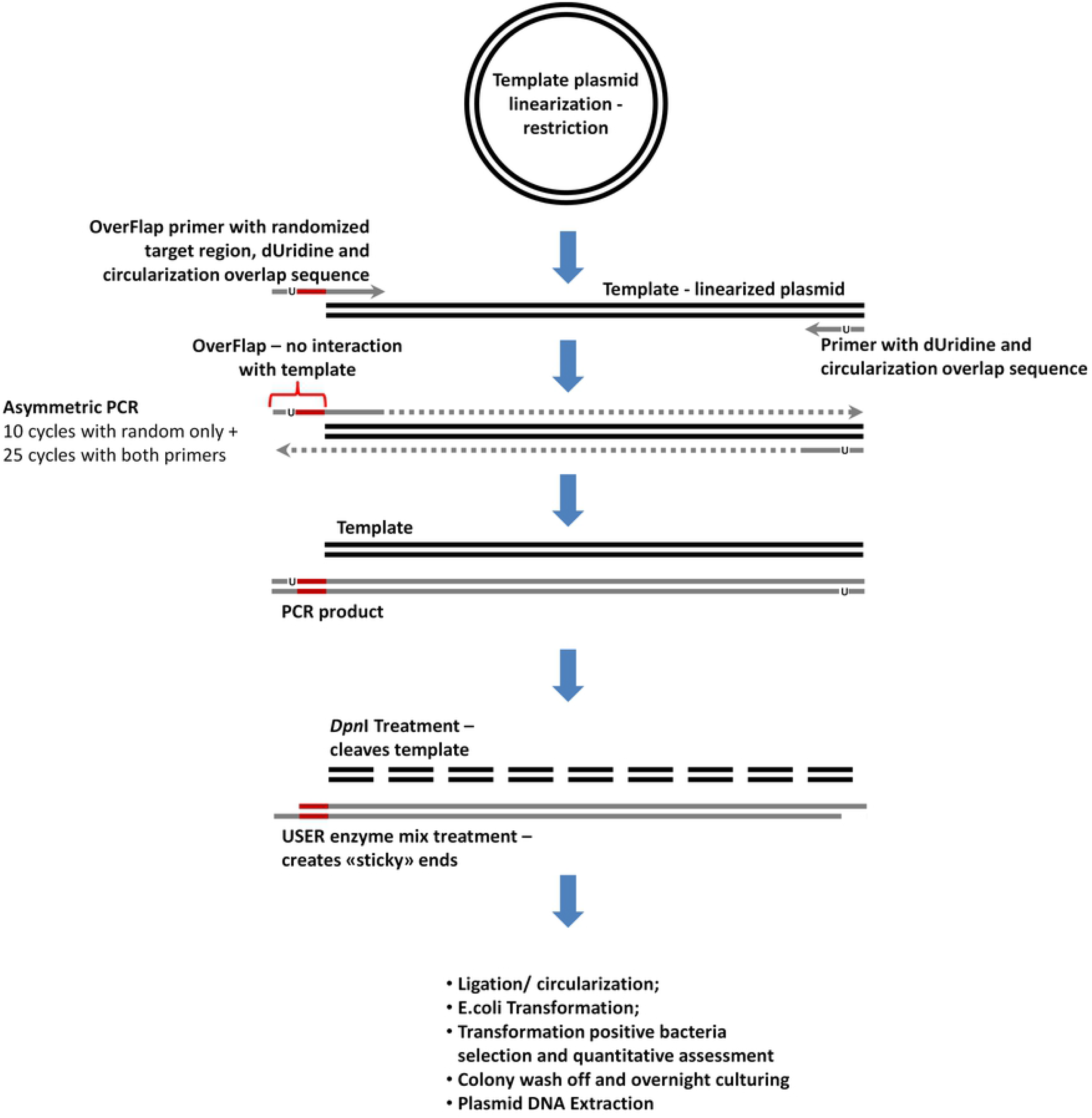
The principal scheme of OverFlap strategy for whole plasmid amplification based randomization of selected plasmid DNA region. The methodological approach following the circularization is identical to the one described in Figure 1, but unlike in previous, here the plasmid is linearized in a way that random region of the longer primer does not interact with template and additional stage of asymmetric PCR with only random region containing primer is performed to increase the number of sequence variants within final library. U represents deoxy uridine within employed primers; Randomized region is represented by red line; The template DNA is represented by black line; The synthesized chain is represented by grey line; Dotted line represents DNA synthesis; and dashed line represents template DNA degradation by DpnI restriction endonuclease.

### Massive parallel sequencing employing IonTorrent PGM system

Thus randomized region plasmid was amplified using 0.05 μg of plasmid DNA, 1x Phusion U Multiplex PCR Master Mix, 10 pmol of NGS-RanSeq-Fw and NGS-RanSeq-Rs-77 or NGS-RanSeq-Rs-69 primers (Table 1) and water to final volume of 10 μl, thermal conditions were as follows: 98°C for 30 sec; followed by 25 cycles of 98 °C - 10 sec, 71.5 °C - 15 sec, 72 °C – 15 sec and finalized with 72 °C - 7 min. Afterwards the repertoire of acquired products was assessed employing agarose gel electrophoresis and in the case of success, which was marked by presence of 270 bp fragment, acquired products were then purified with NucleoMag NGS Clean-Up and Size Select kit (Macherey-Nagel, Germany) according to manufacturer’s instructions. The quality and acquired amount of the amplicons was assessed using Agilent High Sensitivity DNA Chip kit on Agilent 2100 BioAnalyzer (Agilent Technologies, USA) and Qubit dsDNA HS Assay Kit on Qubit 2.0 Fluorimeter (Thermo Fisher Scientific, USA).

Prior to clonal amplification each library was diluted to 12 pM and pooled. The Ion PGMTM Hi-QTM View OT2 kit (Life Technologies, USA) and Ion OneTouch DL instrument (Life Technologies, USA) were used for template generation. The template-positive ISPs (Ion Spheres™ Particles) were enriched using Dynabeads MyOne™ Streptavidin C1 beads (Life Technologies, USA) and Ion OneTouch ES module. ISP enrichment was confirmed using the Qubit 2.0 fluorimeter (Life Technologies, USA). The sequencing was performed on Ion 318 v2 chip and Ion Torrent PGM machine employing the Ion PGMTM Hi-QTM View Sequencing kit (Life Technologies, USA). All procedures were carried out according to manufacturer’s instructions and each run was expected to produce approximately 150,000 reads per sample. Due to que and run availability at genetic analysis facility sequencing runs for WPA and OverFlapWPA were carried out with the target read length of 200 bp while for OverFlapAsymWPA - 400 bp. Following the sequencing procedure, the individual reads were filtered by the PGM software to remove low quality reads. Sequences matching the PGM 3’ adaptor were automatically trimmed. All PGM quality-approved, trimmed and filtered data were exported as fastq files.

### Sequencing data analysis

Following data acquisition cutadapt v1.15 was used to test all acquired reads for the presence of sequences that surrounded the randomized region and to extract randomized region along with the 3 adjacent nucleotides from both sides (last codon of α-factor and stop codon). Further employing Needleman-Wunsch algorithm as implemented in SeqAn 2.4.0 all thus acquired reads were aligned against the template (α-MSH plus 3 adjacent nucleotides from both sides) and for further analyses were retained only those for which the first and last trinucleotides matched the template and lengths were divisible by 3. Remaining reads were divided in two groups: 1) target group of insufficiently randomized sequences, which contained reads that aligned with less than 10 mismatches and 3 gaps, and 2) remaining – randomized sequence reads. Afterwards, since the initial purpose of this study was to create random peptide expression library reads from both groups were translated employing in-house developed script and all unique sequences were quantified.

## RESULTS

As already explained in previous sections the main purpose of this study was to create a plasmid library that would enable the production of random peptides in yeast *S. cerevisiae* expression system. Since pharmacophores for majority of biologically active peptides are below 18 aa we chose to create a plasmid that would be capable of peptide secretion (would contain peptide secretion signal) and would already contain a peptide of similar length (due to our previous research experience we selected α-MSH) whose coding sequence would later on be randomized. For the production of random peptides we selected a p426GPD expression plasmid, which is compatible with S. cerevisiae expression system. The selection of given vector was based on its wide application in yeast expression systems, knowledge that GPD is one of the strongest yeast promoters [60] and recommendations from Dr. Dowell. For the introduction of α-factor secretion signal and α-MSH fusion protein into selected vector we employed PCR based mutagenesis approach, which has been successfully employed in our laboratory on multiple occasions, therefore neither deviations from initial plan nor any unforeseen problems were encountered. The envisioned plasmid was created in first attempt and success was verified by Sanger sequencing (data not shown).

### Randomization employing modified whole plasmid amplification strategy (WPA)

Our initial studies of literature revealed that there are two suitable strategies for our study: the cassette mutagenesis, which relies on restriction/ligation for insertion of randomized sequence containing fragment [51-53] and whole plasmid amplification with randomized sequence containing oligonucleotide [6, 50]. Our rather extensive experience in cloning and expression of various mammalian genes suggests that the ligation of two fragments is less efficient than circularization, because un like the latest it is a two stage process, where the first stage requires joining of two spatially unrestricted DNA ends, while the second in essence is circularization of newly formed molecule through joining of both ends which now are spatially restricted to relatively proximal location. Therefore to gain a greater number of clones with randomized sequence and possibly a higher level of diversity in randomized sequences we chose to employ the whole plasmid amplification and following circularization approach. However, although this approach in our laboratory has been used on several occasions, because of the recombination based end joining, we found it to be less reliable than more traditional overlapping end PCR based mutagenesis, which in its final stage involves employment restriction endonucleases and creation of more reliable DNA “sticky ends”. Thus it was decided that selected strategy shall be augmented with either introduction of restriction site after the randomized region or employment of deoxy uridine containing primers to create “sticky ends” after the treatment with USER enzyme mix (contains Uracil DNA glycosylase (UDG) and Endonuclease VIII) [9]. Since the last one allows creation of longer overhangs that in theory should increase the efficiency of circularization, it was the method of choice for our further activities.

One of the first activities that were undertaken within the scope of randomization was design of the primers. Our strategy here was based on employment of one long reverse strand oligonucleotide, which at its 3’ end contains sequence that is complementary 18 nucleotides of α-factor secretion signal CDS’s 3’ end, followed by 54 random nucleotides, stop codon and additional 9 nucleotides where the first thymidine was replaced with deoxy uridine, and one short forward strand oligonucleotide, which is at its 3’ end contains sequence that is complementary 23 reverse strand nucleotides of p426GPD plasmid that follow immediately after stop codon of α-MSH CDS, these are followed by additional 9 nucleotides that are complementary to reverse primer’s 5’ nucleotides and also here the first thymidine was replaced with deoxy uridine. (Table 1)

For the amplification of whole plasmid we selected Phusion U Multiplex PCR Master Mix, because it is optimized for amplification of difficult targets and more importantly it contains high fidelity polymerase (Phusion U) which tolerates presence of Uridine within template and growing DNA chains. The amplification itself was successful and there was no DNA degradation neither prior no after treatment with USER enzyme mix and *Dpn*I restriction endonuclease, which specifically cleaves only methylated template plasmid DNA (Figure 3). Therefore acquired DNA fragment was circularized, transformed in competent cells and seed onto a petri dish. The next day assessment of colony forming units by OpenCFU software revealed that there are ∼3 802 colonies on our 8.8 mm petri dish (Figure 4a). However it should be noted that the actual numbed could be greater by as much as 20%, because we observed that resolution of acquired images (1733×1733 pixels – maximum for our equipment) was insufficient for software to reliably identify smaller colonies and distinguish individual ones within dense colony clusters, but despite this drawback OpenCFU provided a simple, reliable and unbiased colony estimation for monitoring of case to case transformation efficiency. The colony number that was acquired from this experiment was nevertheless, lower than expected and insufficient to gain a well-represented random sequence library of 54 nucleotide length, but is was sufficient for assessment of randomization process efficiency, therefore colonies were washed off and acquired cell culture inoculated in liquid media for overnight growth and plasmid extraction.

**Figure 3.**
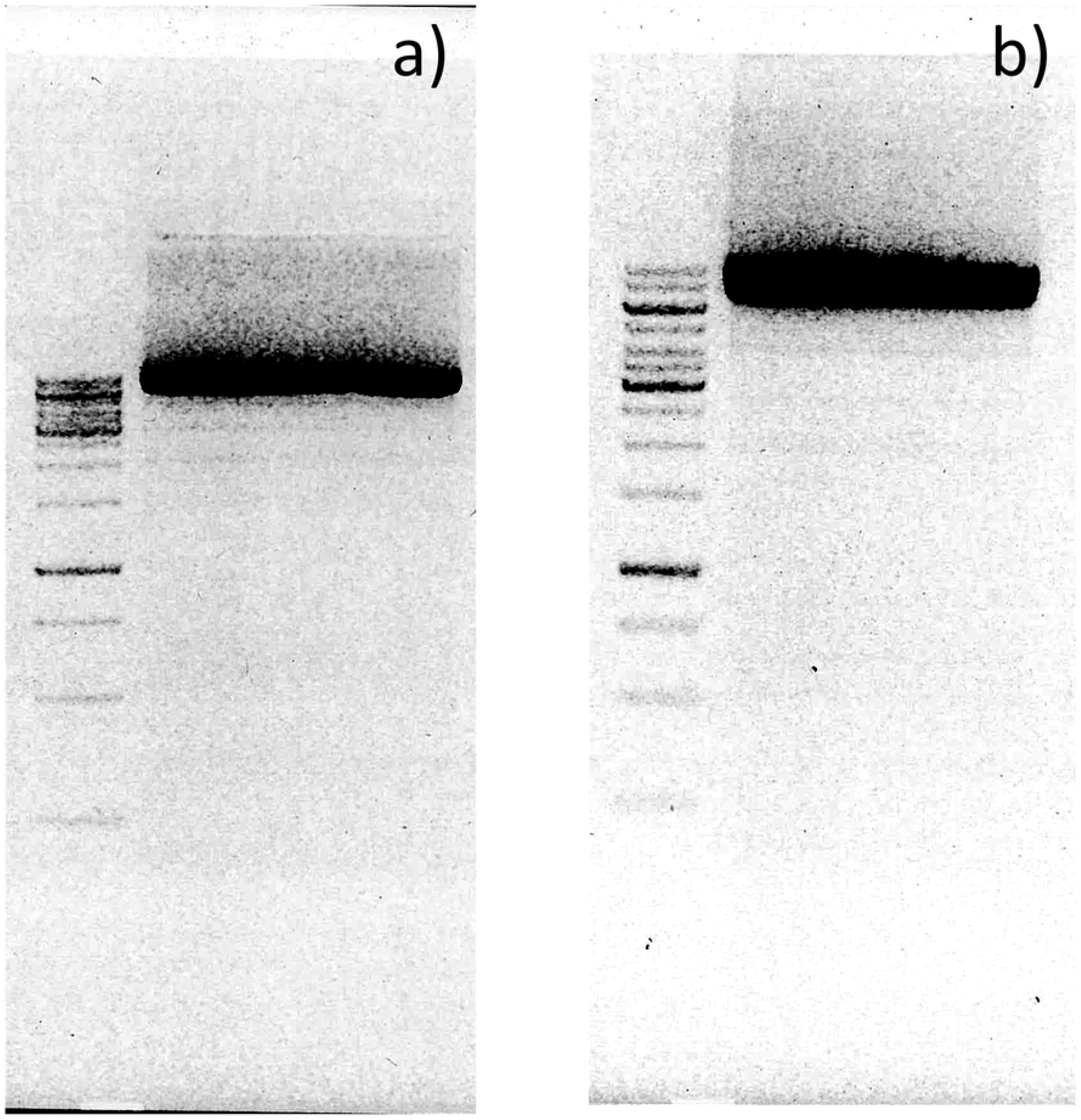
Visualization of whole plasmid amplification (WPA) products in agarose gel electrophoresis a) prior *Dpn*I and USER enzyme mix treatment and b) after treatment indicates that whole plasmid amplification process was successful and the size of amplification product was not affected by treatment i.e. no degradation was observed. First line in each gel contains 2 μl of GeneRuler 1 kb DNA Ladder (Thermo Fisher Scientific, Lithuania), the uppermost band is 10 000 bp and three brightest bands are 6 000 bp, 3 000 bp and 1 000 bp. The second lane in each gel contains 5 μl sample of acquired reaction products.

**Figure 4.**
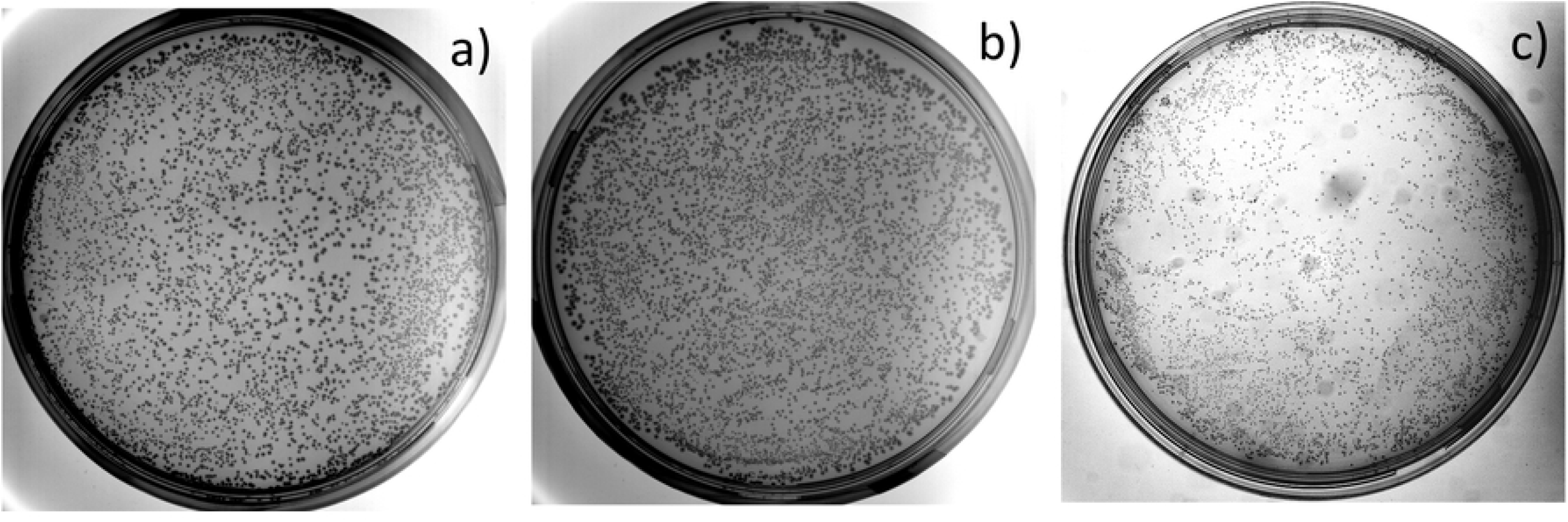
Photographic pictures of 8.8 mm diameter petri dish with selective Amp+ media containing colonies of *E*.*coli* Dh5α strain transformed with circularized product of a) whole circular plasmid amplification (WPA) - 3 802 colonies; b) OverFlapPCR based whole plasmid amplification (OverFlapWPA) - 4 534 colonies; and c) OverFlapPCR based asymmetric whole plasmid amplification (OverFlapAsymWPA) - 4 865 colonies. All procedures were done as described in Materials and methods section.

Our initial attempt for assessment of randomization process efficiency involved Sanger sequencing and acquired results confirmed that randomization as such was successful (Figure 5), although the major peak could be observed at every position, but they differed between sequenced strands rendering the assessment as inconclusive. Therefore, to gain an in depths understanding on sequence composition of created randomized plasmid library we performed NGS sequencing employing IonTorrent PGM sequencing technology.

**Figure 5.**
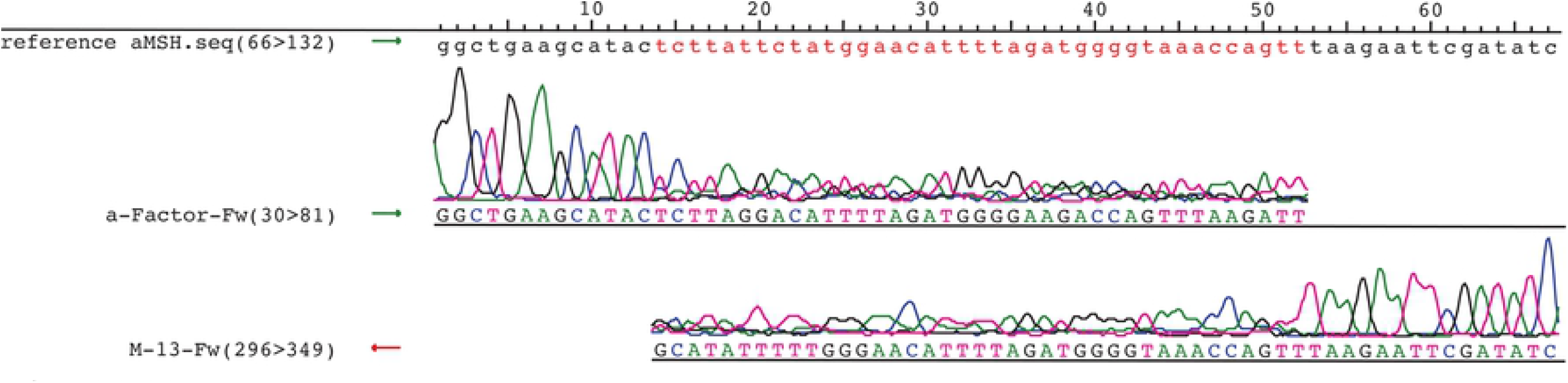
Assembly of WPA plasmid Sanger sequencing capillary electrophoresis chromatograms. Sequencing was performed employing a-Factor-Fw primer for forward strand sequencing and M13-Fw primer for reverse strand sequencing. The fragment containing sequence of template - α-MSH (red font) was used as reference for this analysis.

The preparation of IonTorrent PGM compatible sequencing libraries was performed in similar manner as libraries for microbial 16S rRNA community analysis [61] – i.e. the primers that included both target sequence and IonTorrent technical sequences (adapters and barcodes) were used for amplification of randomized region. The size of target amplicon was 273 bp (vs 54 bp of randomized region), because employed primers were targeted at sequences that were located 98 bp upstream and 62 bp downstream of randomized region, to both provide reliable reference for identification of randomized region at data analysis stage and enable reliable size separation from primer dimers that form during any PCR based amplification. Since it is a requirement of all NGS technologies the quality of acquired sequencing library was assessed employing capillary electrophoresis and the result revealed that contrary to expected our library was formed by products of various sizes, which indicated that in addition to randomization of target sequence, some deletions and duplications have also occurred (Figure 6 WPA) and subsequently acquired sequencing data partially confirmed this suspicion, because many of randomized translated sequences without template bias were shorter than 18 aa (54 nucleotides). The data also revealed that ∼50% of acquired reads were highly similar to template, confirming the concerns of researchers that any template based randomization leads to considerable bias towards the sequence of the template, which in essence jeopardizes any attempts for acquisition of random sequence library without template bias. In addition the diversity of randomized sequences was also lower than expected, because 969 unique protein coding sequences (vs ∼3 802 colonies) were identified and only 4 of these were encoded by ∼60% of the reads, even 4 of protein coding sequences that resembled template were encoded by ∼49% of the reads. Also, according to acquired data, the average frequency of specific aa occurrence at every position did not resemble theoretical frequency of aa occurrence that should be observed in the case of truly randomized peptide coding library (Table 2, Supplementary table: sheet 1 and sheet 2). It should also be noted that the true number of unique protein coding sequences is probably even lover due to the PCR based library preparation and IonTorrent technology sequencing artefacts, in fact the latest is renowned for its problems with base calling in homo polymers.

**Table 2.** General characteristics of acquired sequencing data and an overview of results that were acquired during following data analysis. WPA represents randomization through whole circular plasmid amplification; OverFlapWPA – OverFlapPCR based whole plasmid amplification; OverFlapAsymWPA - OverFlapPCR based asymmetric whole plasmid amplification.

|  | WPA | OverFlapWPA | OverFlapAsymWPA |
| --- | --- | --- | --- |
| **Number of observed colonies** | **3 802** | **4 534** | **4 865** |
| **Number of Sequencing reads** | **140 960** | **139 461** | **806 671** |
| **Mean Read Length** | **147 bp** | **141 bp** | **163 bp** |
| **Number of target region sequence reads (passed quality filters)** | **62 636 (44.44%)** | **94 609 (67.84%)** | **611 478 (75.8%)** |
| Number of reads that encode peptides with COP $\geq 1$ | 60 977 | 87 791 | 103 747 |
| Number of reads that encode peptides with COP <1 | 1 659 | 6 818 | 507 731 |
| **Number of unique protein coding sequences** | **969** | **4 636** | **51 504** |
| <sup>1</sup>Number of unique protein coding sequences with COP $\geq 1$ | 84 | 464 | 627 |
| Number of unique protein coding sequences with COP <1 | 885 | 4 172 | 50 877 |
| **COP of all unique protein coding sequences** | **3 802.00** | **4 534.00** | **4 865.00** |
| COP of all unique protein coding sequences with individual COP $\geq 1$ | 3 701.30 | 4 207.25 | 825.42 |
| <sup>2</sup>COP of all unique protein coding sequences with individual COP <1 | 100.70 | 326.75 | 4 039.58 |
| **Number of insufficiently randomized reads (% of total)** | **31 447 (50.21%)** | **3 (0.003%)** | **0 (0%)** |
| Number of insufficiently randomized reads that encode peptides with COP $\geq 1$ | 31 110 | 0 | 0 |
| Number of insufficiently randomized reads that encode peptides with COP <1 | 337 | 3 | 0 |
| **Number of unique insufficiently randomized protein coding sequences** | **117** | **1** | **0** |
| <sup>3</sup>Number of unique insufficiently randomized protein coding sequences with COP $\geq 1$ | 9 | 0 | 0 |
| Number of unique insufficiently randomized protein coding sequences with COP <1 | 108 | 1 | 0 |
| **COP of all unique insufficiently randomized protein coding sequences** | **1 908.83** | **0.15** | **0.00** |
| COP of all unique insufficiently randomized protein coding sequences with individual COP $\geq 1$ | 1 888.37 | 0.00 | 0.00 |
| <sup>4</sup>COP of all unique insufficiently randomized protein coding sequences with individual COP <1 | 20.46 | 0.15 | 0.00 |
| **Number of randomized reads (% of total)** | **31 189 (49.79%)** | **94 606 (99.997%)** | **611 478 (100%)** |
| Number of randomized reads that encode peptides with COP $\geq 1$ | 29 867 | 87 791 | 103 747 |
| Number of randomized reads that encode peptides with COP <1 | 1 322 | 6 815 | 507 731 |
| **Number of unique randomized protein coding sequences** | **852** | **4 635** | **51 504** |
| <sup>5</sup>Number of unique randomized protein coding sequences with COP $\geq 1$ | 75 | 464 | 627 |
| Number of unique randomized protein coding sequences with COP <1 | 777 | 4 171 | 50 877 |
| **COP of all unique randomized protein coding sequences** | **1 893.17** | **4 533.86** | **4 865.00** |
| COP of all unique randomized protein coding sequences with individual COP $\geq 1$ | 1 812.92 | 4 207.26 | 825.42 |
| <sup>6</sup>COP of all unique randomized protein coding sequences with individual COP <1 | 80.25 | 326.60 | 4 039.58 |
| <sup>1+2</sup>**Number of unique protein coding sequences within libraries** | **185** | **791** | **4667** |
| <sup>3+4</sup>Number of unique insufficiently randomized protein coding sequences within libraries (% of total colonies) | 30 (50.21%) | 0 (0%) | 0 (0%) |
| <sup>5+6</sup>Number of unique randomized protein coding sequences within libraries (% of total colonies) | 155 (49.79%) | 791 (100%) | 4667 (100%) |

\* Expected minimal abundance in %, calculated as 100% divided by number of observed colonies on petri dish

**Figure 6.**
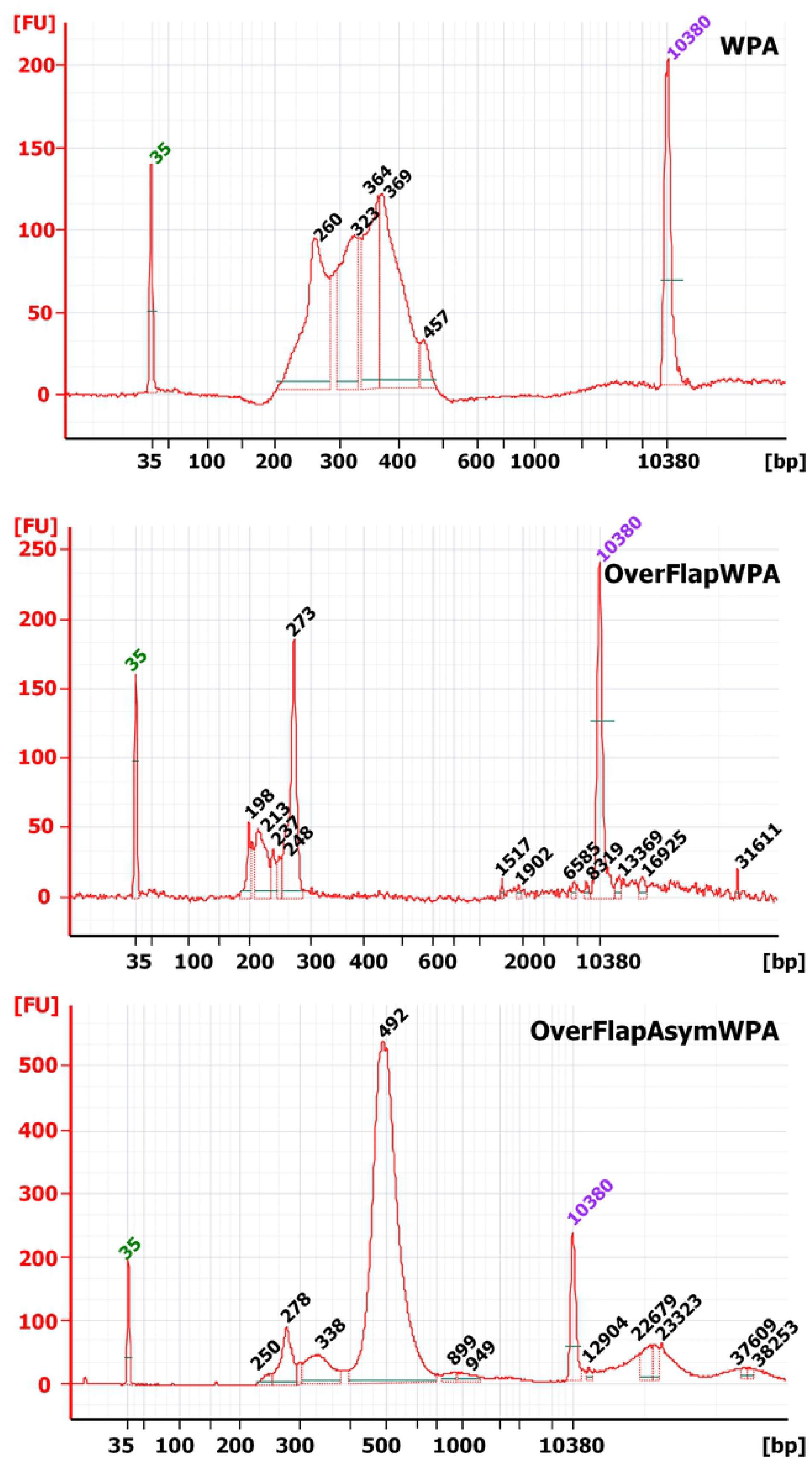
Chromatograms of sequencing library fragment sizing analysis that was carried out on Agilent Bioanalyzer 2100. WPA - whole circular plasmid amplification; OverFlapWPA - OverFlapPCR based whole plasmid amplification; and OverFlapAsymWPA - OverFlapPCR based asymmetric whole plasmid amplification. The 35 bp (green) and 10380 bp (purple) peaks represent Upper and Lower markers that are employed by data analysis software for internal calibration of each individual analysis.

The typical approach that many researchers use to deal with these false reads is setting the requirement for identification of minimal number of identical reads before specific sequence is considered as true [53]. However this approach has several drawbacks: 1) it is rather mechanistic and it is rarely based on some careful considerations; 2) it can be easily bypassed if larger amount of sequencing data is acquired, because the number of false positive reads increases proportionally. Therefore we decided to employ an alternative - an occurrence probability based strategy for assessment of acquired variant number.

As of first, since we were aware of the number of colonies that formed the base of our random plasmid library and it is a well-known fact that transformation of individual bacterial cells with more than one variant is a rare event [62], it was possible to calculate the cumulative occurrence probability (COP) for each of observed variants. In essence this calculation involved multiplication of observed colony count with is abundancy fraction for specific variant, and the later one can be considered as probability of each variant occurrence. Following this, whole variant repertoire was divided in two groups – those for which the COP value was equal or above 1 and those with COP below 1. Thus the first group comprised variants that would certainly occur in our library while the second included those for which the occurrence was not certain. Then we calculated the COP value of each group by summing the COP values of each individual variant, thus acquiring the rough assessment of the number of colonies that in theory would have been occupied by members of each group. The number of variants that was present in our library was calculated by summing the number of variants within the first group, which is the actual number of variants within this group, with the COP value of the second group rounded to the nearest whole number, which corresponds to the approximate number of colonies that were occupied by members of this group and, since predominant majority of bacterial cells are transformed by single variant, it is closer to the actual number of variants in the second group.

Thus after application of these calculations we concluded that WPA approach has resulted in creation of only 185 unique protein coding sequences (Table 2). Although this approach can be considered as complicated, significantly affected by colony estimation accuracy and differences in replication speed of various plasmid variants during bacterial culturing in liquid media, it has several advantages 1) it can be calculated and individually applied to each data set and 2) being frequency based it is less sensitive to fluctuations of sequencing data amount.

### Randomization employing OverFlapPCR based whole plasmid amplification strategy (OverFlapWPA)

As mentioned previously the only proposed solutions for mitigation of template bias problem were adjustment of annealing temperature and Loop-out/loop-in excision of target sequence from the template, but since we intended to 54 bp region the temperature adjustment would not be a feasible option, while by excision of α-MSH CDS we would simply switch the template to sequence that is located downstream of α-MSH CDS, we believed that none of these approaches would solve the issue. Therefore we devised a novel strategy where the prior PCR based whole plasmid amplification template is linearized at the position that is located just before, just after, short distance upstream or short distance downstream of intended randomization site, thus ensuring that template ends with randomization primer’s target site (part that has perfect complementary to template) and during whole plasmid amplification randomized part forms an overhang that sort of “freely flaps” over the end of the template (Figure 2) having minimal interaction with it and mitigating the bias effect, hence we named this approach an OverFlapPCR or OverFlapWPA.

In our case we were able to identify restriction site that was a conveniently located 21 bp upstream of α-MSH CDS within CDS of α- secretion factor. All other activities were carried out identically as in the case of WPA. Just like previously the OpenCFU estimated colony number - 4 534, although a little higher, was still lower than necessary for acquisition of well represented random sequence library, but sufficient to evaluate its diversity. Sanger sequencing of acquired plasmid library (data not shown) confirmed that randomization per se has happened and therefore we proceeded with generation sequencing library and its quality assessment. Data from Bioanalyzer revealed that the proportion of shorter and longer fragments was significantly decreased and single peak of target 273 bp could be clearly identified. (Figure 6 OverFlapWPA) NGS analysis revealed that in comparison with traditional WPA the number of unique protein coding sequences was increased more than 4 times reaching 4 636 and what was even more important the overwhelming majority reads (99.997%) were randomized, bearing no or little similarity to CDS of α-MSH. In addition the abundancy of most represented sequence was below 5% and the total abundancy of four most represented decreased to only ∼8.2%. Also, although the application of previously presented COP based calculations significantly decreased the number of unique reads to 791, it was still more than 4 times higher than that of WPA. Also the average frequency of specific aa occurrence at every position displayed a greater similarity to theoretical frequency of aa occurrence in the case of truly randomized peptide coding library, thus highlighting that significant imprrovements were achieved. (Table 2, Supplementary table: sheet 1 and sheet 2) Taken together these results clearly demonstrate that sequence diversity within OverFlapWPA generated libraries is significantly greater than within those that were generated by traditional WPA approach.

### Randomization employing asymmetric OverFlapPCR based whole plasmid amplification strategy (OverFlapAsymWPA)

Never the less, despite the previously described success we believed that, due to the fact that ∼8.2% reads encoded four most represented decreased, the sequence diversity within acquired libraries could still be improved. In our understanding the explanation for this phenomenon might be related to peculiarities of PCR, where those sequences that are generated during the first cycles are amplified at greater speed than others. In our view there were two solutions to this problem. 1) to increase the template concentration which is accompanied by decrease of amplification cycles or 2) perform several cycles of asymmetric PCR with randomization primer to linearly increase the concentration of already randomized templates without introduction of early cycle amplification bias. Since the employment of the first option might increase the template background within randomized library we decided to proceed with the second. Thus we altered the first 10 cycles of PCR procedure by carrying it out with only randomization primer, while the remainder of procedures was identical to previous ones. These activities resulted plasmid library that was acquired from 4 865 transformation positive colonies. As previously the success of randomization procedure per se was confirmed by Sanger sequencing (data not shown). Peculiar in this case were the results of agarose gel electrophoresis and capillary electrophoresis based DNA fragment analysis. They revealed that the size of the major library PCR product was 492 bp while the amount of expected 273 bp fragments, although shifted by 5 bp, was significantly lower (Figure 6 OverFlapAsymWPA). Therefore it was decided to perform an additional purification with size selection beads to remove any DNA fragment larger than 300 bp, but the analysis of acquired DNA sample revealed the same pattern – major peak was larger than expected. Our further attempts with ∼250bp band excision from agarose and following DNA purification returned same results. Therefore it was concluded that the apparently the larger fragments are actually some kind of “products” of our target fragment. Our best speculation on this matter is that due to high level of randomization and PCR related cyclic temperature induced denaturation-renaturation, there is a significant proportion of double stranded DNA molecules within our sequencing library for which both strands are not perfectly matched leaving these unmatched bases free for interaction with other DNA molecules and formation of something that is akin to G-quadruplexes [63], but, since these interactions are due to random nature of library sequence and not the result of intelligent design or evolutionary selection, they are not stable and it is possible that these quadruplexes are constantly forming and disbanding during DNA travel through electrophoresis polymer, which might result in shift of the peak/band size towards something that is smaller than the sum of both molecules. Be as it is, our numerous purifications resulted in significant loss of sequencing library, therefore it was recreated purified and sequenced as described in materials and methods section. Acquired sequencing data also confirmed that the input library was of correct size, because despite of the coincidental sequencing of 400 bp long reads (performed due to requirements of other libraries that were sequenced during specific sequencing run) the mean read length was only 163 bp. It should also be noted that this sequencing run resulted in acquisition of significantly larger amount of data (806 671 reads) than the previous runs (∼140 000 reads), thus all abundancy and diversity results should be considered in context with this fact. Sequencing data analysis revealed that in comparison with OverFlapWPA the number of unique protein coding sequences was 11 times higher reaching 51 504, which is considerable increase because it resulted from data amount that was ∼only 6.5 times larger. Interestingly in spite of the increased data amount all of the reads were randomized bearing no or little similarity to CDS of α-MSH. Even more, the abundancy of most represented sequence was below 0.1% and the total abundancy of four most represented was below 0.25%. Also, although the application COP based calculations significantly decreased the number of unique reads to 4 667, it was still almost 6 times higher than that of OverFlapWPA. Similarly further improvements were also observed in calculations of average frequency of specific aa occurrence at every position, here experimentally acquired numbers displayed even greater resemblance to theoretical frequencies of aa occurrence within of truly randomized peptide coding library (Table 2, Supplementary table: sheet 1 and sheet 2) Thus it is clear that introduction of asymmetric PCR stage resulted in acquisition of random plasmid library with even greater sequence diversity.

## DISCUSSION

Recent developments in process automatization and high throughout data acquisition technologies have started a new era in many research fields of life and medical sciences. Being at the frontline of known and trying to develop something that cannot be found or has not been observed in nature, through series of creation and effect observation experiments, the fields of protein engineering and synthetic biology have also greatly benefited from introduction of these technologies in their everyday practices. Introduction of NGS technologies has been particularly beneficial to disciplines that involve studies or improvement of proteins through their coding sequence randomization. Although, as it was explained in introduction section, there are multiple methodologies for randomization of target sequences that fit various research strategies, currently technologically most challenging is reliable randomization of protein segments that are larger 9 bp, because strategy involving cassette replacement require convenient localization of two research sites and due inefficiency of two fragment ligation are able to produce relatively low number of transformation positive colonies, while, as it was also clearly demonstrated in this study, PCR based whole plasmid amplification suffers from significant randomization bias towards the template sequence.

The OverFlap PCR strategy that we present in this study provides a simple and reliable solution to said sequence bias problem. As it was demonstrated in our OverFlap whole plasmid amplification (OverFlapWPA) experiment the linearization of the plasmid at the position that is adjacent to randomization site prevents primer’s random sequence containing segment interaction with template during the amplification, which results in acquisition of template unbiased sequence library. Even more, our asymmetric OverFlapWPA (OverFlapAsymWPA) experiments demonstrate that even higher sequence diversity can be achieved if principles of asymmetric PCR are applied during the first cycles of OverFlap whole plasmid amplification. An additional problem that we encountered during our initial WPA experiments was high prevalence of reads that were shorter than randomized region (Figure 6 WPA, Supplementary table: sheet 1). Although we cannot provide any reasonable explanation to this phenomenon beyond mismatch induced degradation by either polymerase or *E*.*coli* repair machinery, it seems that introduction of asymmetric amplification has resolved the issue (Figure 7).

**Figure 7.**
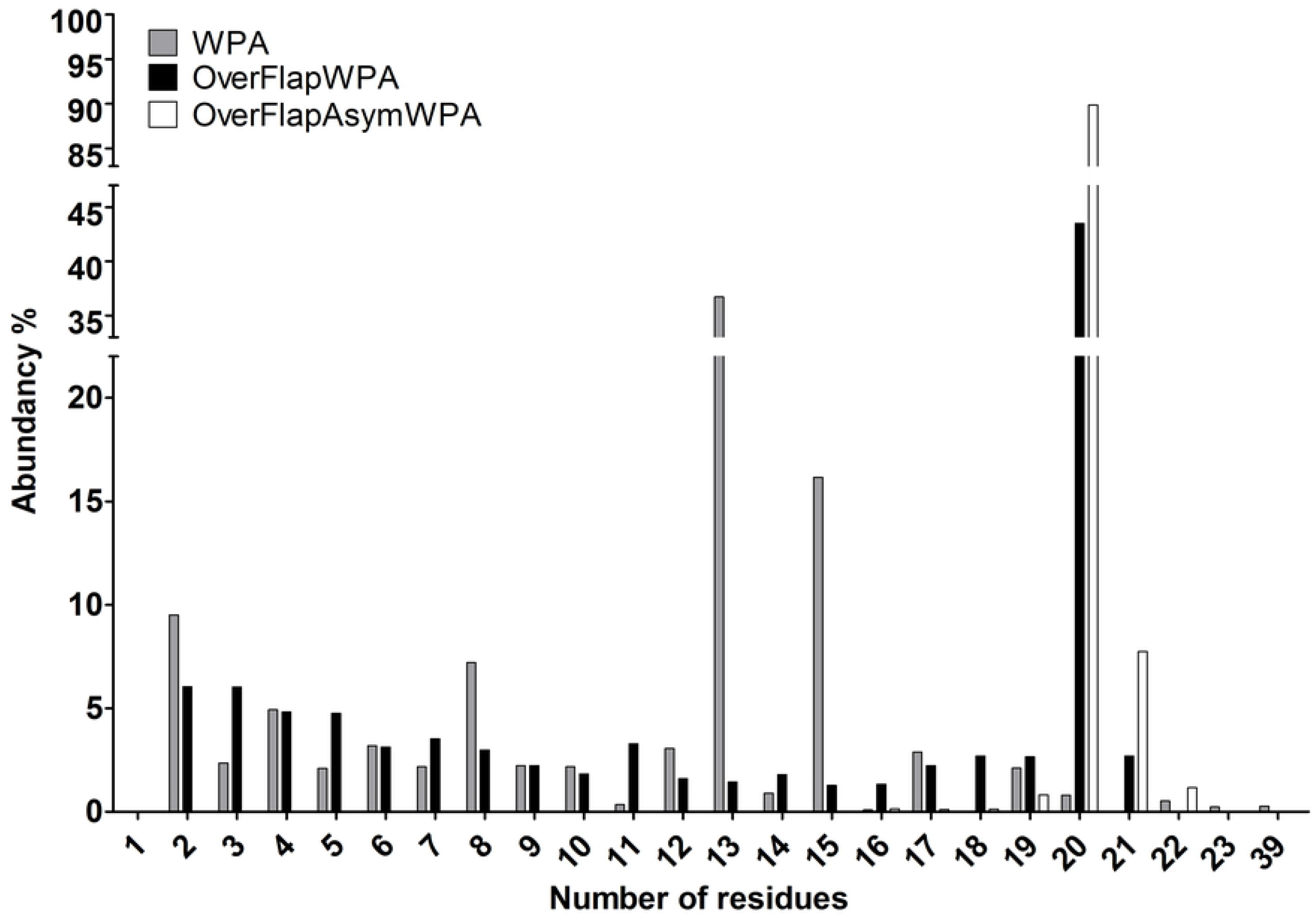
Quantification of randomized region encoded peptide according to size. As described in materials and methods section data includes last aa of α-factor secretion signal CDS and stop codon, thus the reals size of. In the case of 1 last aa of α-factor secretion signal CDS was missing.

Although it is very effective for acquisition of randomized library without template bias, the described methodology has several drawbacks.

As of first the number of acquired transformation positive colonies is far too low to ensure that all possible variants are be represented in acquired library of larger scale randomization. According to very rough calculation on the average we were able to acquire ∼4 400 colonies per transformation, thus under ideal conditions, where all possible sequence variants are evenly represented, this number shall be sufficient to reliably randomize no more than two codons (46=4 096). Off course, this situation would be less of the problem for those researchers that are interested in identification of novel biologically active peptides, because having library 4 000 reliably random peptides is better than having library of 4 000 peptides with significant bias towards template, but for those that wish to randomize specific section of known protein during single experiment this might indeed be a significant drawback. In our opinion there are two types of solutions to this problem. The first would involve performance of repeated randomizations and transformation positive colony collections until the desired number of colonies or the level of variations determined by NGS is reached, while the second would involve employment of entirely different circularization strategy. In our opinion the best alternative would be Gibson assembly, because several authors have reported acquisition of up to 108 of colony forming units [49]. An additional advantage of Gibson assembly would be the correction of possible mismatches between strands, because, due to close proximity to the end of DNA fragment, randomized region for one of the strands shall be destroyed and rebuilt anew using remaining strand as template.

As of second the reliance on restriction for linearization of template limits the choice of randomization sites. The workarounds in this case would be either introduction of restriction sites within template through employment of site directed mutagenesis or linearization through whole plasmid amplification. The choice of introduced restriction site in the first case would mater little, as long as it would be unique within the chosen template, because during randomization process altered nucleotides can be reverted to desired state. The advantage of restriction enzyme approach is that after the randomization *Dpn*I restriction enzyme can be used for reliable fragmentation/destruction of template DNA, while the disadvantage is the necessity to perform additional mutagenesis. The advantage of the whole plasmid amplification on the other hand is that plasmid can be linearized at virtually any site within single step, but due to lack of methylation such template shall be chemically indistinguishable from newly created strands and selective digestion by *Dpn*I restriction enzyme won’t be possible. Nevertheless we believe that due to its simplicity the whole plasmid amplification approach would be more attractive than introduction of restriction site and we currently see two workarounds for mitigation of the template background issue. The simplest one is to exclude the intended overlap and randomization target regions from linearized template through employment of PCR primers that anneal to flanking regions of the randomization target and are directed away from it, thus lack of complementary region shall significantly decrease the probability of circularization while in the case of Gibson assembly end joining shall be nearly impossible. The other workaround would be to use modified dNTP mix, where dTTP is replaced with dUTP in combination with uridine tolerant high fidelity polymerase for template amplification (such as dNTP/dUTP Mix and Phusion U Hot Start DNA Polymerase (Thermo Fisher Scientific, USA)). In this case addition of previously described USER enzyme mix to post-randomization mixture shall result in cleavage and elimination of created linear template DNA in a manner that is similar to *Dpn*I cleavage of methylated sites. Also both of the mentioned workarounds are not mutually exclusive and can be employed in combination.

Taking together the information that has been presented thus far on the subject of possible improvements of our developed randomization methodology, we propose and recommend following workflow for our readers.

The linearization of template should be performed employing PCR based whole plasmid amplification. Here, as an option, the employment of reaction mix with dUTP instead of dTTP and uridine tolerant high fidelity polymerase is applicable. The primers should have perfect complementarity to template and should be designed to exclude both randomization target region and circularization overlap region from linearized randomization template. Acquired product should be treated with *Dpn*I and purified preferably with some size selection methodology (agarose gel excision or magnetic beads) to exclude remaining fragments of template, any nonspecific PCR products and primer dimers. Acquired product should then be used as template for randomization through Asymmetric OverFlap Whole Plasmid Amplification, where first 10 or more PCR cycles are performed employing only randomization primer that also contains overlap sequence and additional 25 cycles are performed employing both primers. If dUTP was used then at this stage acquired products should be treated with USER enzyme mix and purified if not then just purified. Thus acquired randomized liner DNA should be circularized employing Gibson assembly and transformed in competent *E*.*coli* cells of suitable strain. Cells should then be seeded on petri dish to assess the number of transformation positive colonies and thus the theoretical maximal diversity of created library. After that colonies should be washed off and inoculated in liquid media for additional propagation prior plasmid extraction. The assessment of randomization success and diversity of acquired library should be performed employing suitable sequencing technology. The randomization procedure should be repeated each time new plasmid library stock is needed. However if frozen stocks are created then it should be stored in aliquots of sufficiently large bacterial cell number to maintain acquired diversity of random plasmid library and whole volume of single the aliquot should be used for inoculation. Principal scheme of the procedure is presented in Figure 8.

**Figure 8.**
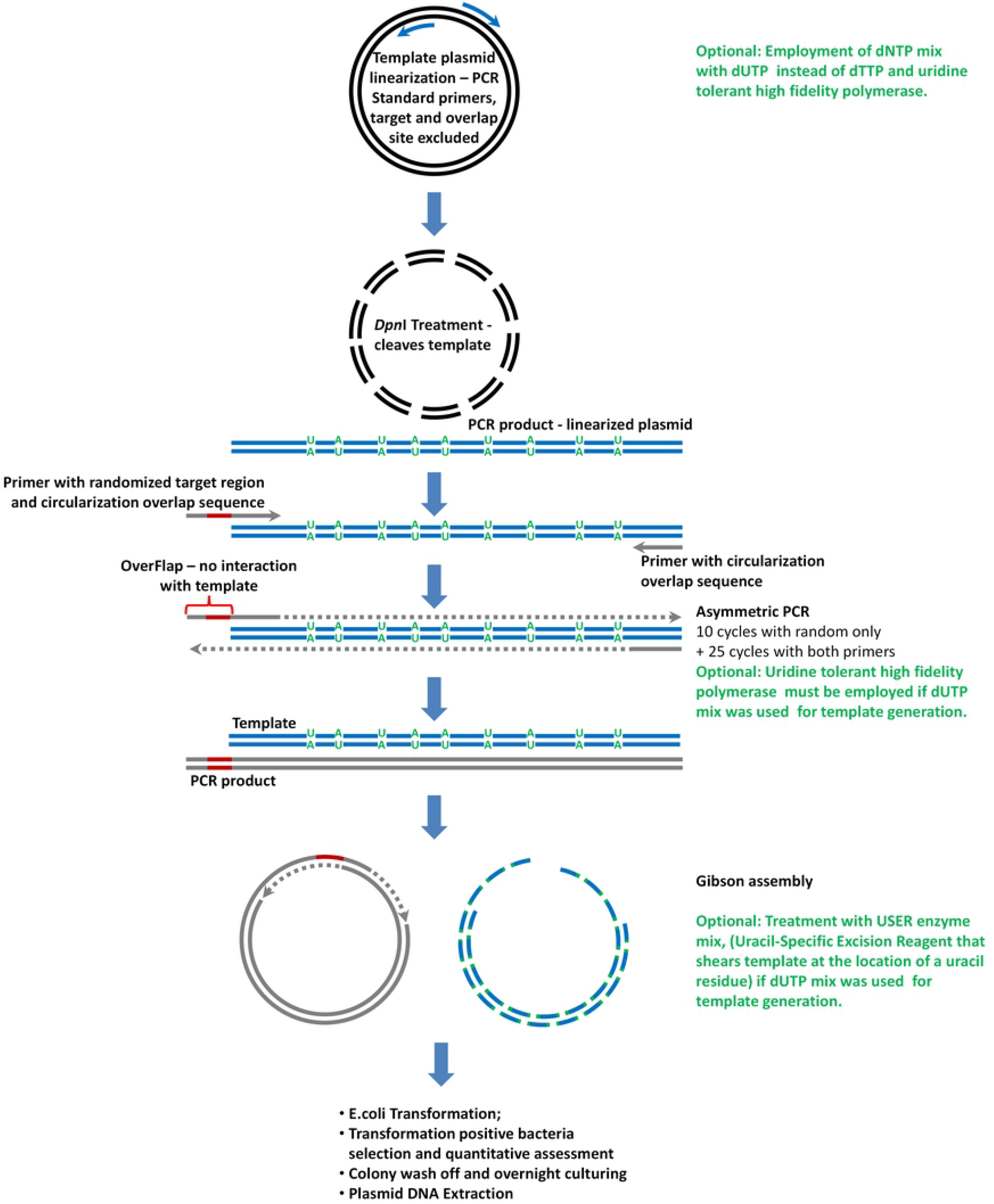
The principal scheme of proposed future OverFlap whole plasmid amplification based randomization of selected plasmid DNA region. Unlike in current iteration here the plasmid is linearized employing whole plasmid amplification and circularization is performed employing Gibson assembly. As an optional strategy for control of unwanted template circularization, one can use 1) dNTP mix with dUTP instead of dTTP and uridine tolerant high fidelity polymerase during template linearization to generate uridine containing template, 2) uridine tolerant high fidelity polymerase during asymmetric PCR and 3) USER enzyme mix (Uracil-Specific Excision Reagent) to selectively cleave template DNA. Randomized region is represented by red line; the template DNA is represented by black line; the synthesized chain is represented by grey line; Dotted line represents DNA synthesis.

Although the methodology presented here is directed towards employment of whole plasmid amplification strategy, the principle of not having template at the randomization site can also be used in other randomization strategies. Thus, for example, in any probe based randomization instead of employment of single and large template, the usage of fragmented one with gaps at selected randomization sites should be considered. Even more since Gibson assembly can be applied for creation of single construct from multiple fragments [64], our OverFlap PCR approach can also be employed for randomization of multiple sites.

In summary with this article we present a novel approach for reliable introduction of random and unbiased nucleotide sequence within virtually any site of the plasmid which we named the “OverFlap PCR” to emphasize the distinction from “Overhang PCR” and “Overlap PCR”. The method is based on employment of randomised region containing primer and linearized plasmid template in partially asymmetric whole plasmid amplification, which is followed by circularization. The linearization of plasmid is carried out in a manner that prevents or minimizes primer’s randomized region interaction with the template. Massive parallel sequencing on IonTorrent PGM machine confirmed that acquired library is random and displays no template bias. At the end of the article we also discuss the drawbacks of developed methodology and present plan for future improvements.

## ACKNOWLEDGEMENT

We would like to thank Dr. Simon Dowell for provision of necessary plasmids and aid in selection of plasmid with appropriate promoter and Ms. Linda Lazdiņa for her input in modification of source plasmid.

## AVAILABILITY

cutadapt v1.15 is software tool that removes adapter sequences from sequencing read. It is available in the GitHub repository (https://github.com/marcelm/cutadapt/).

SeqAn 2.4.0 is an open source C++ library of efficient algorithms and data structures for the analysis of sequences with the focus on biological data. It is available in the GitHub repository (https://github.com/seqan/seqan/).

OpenCFU is a free software that should facilitate (and render more reproducible) the enumeration of colony forming unit (CFU). It is available in the SourceForge repository (http://opencfu.sourceforge.net/).

## FUNDING

This work was supported by European Regional Development Fund (ERDF) [Project No.: 1.1.1.1/16/A/055];

## CONFLICT OF INTEREST

No conflict of interest is identified.

## Notes

### Competing Interest Statement

The authors have declared no competing interest.

